# Perseus: Lineage-Aware Refinement of Kraken2 Taxonomic Classification for Long Read Metagenomes

**DOI:** 10.64898/2026.03.06.710148

**Authors:** Matthew H. Nguyen, Michael C. Schatz

## Abstract

**Motivation:** Long-read metagenomic sequencing improves assembly contiguity and enables genome-resolved analysis of complex microbial communities, but accurate taxonomic classification of long reads and assembled contigs remains challenging. Highly scalable k-mer-based classifiers such as Kraken2 frequently over-assign fine-rank taxonomic labels when applied to long-read data, producing high false positive classification rates driven by sparse or localized k-mer matches, particularly in microbiomes with extensive taxonomic novelty.

**Results:** We present **Perseus**, a lineage-aware confidence estimation framework for taxonomic classification that models the spatial distribution and hierarchical consistency of k-mer evidence along sequences. This formulation reframes taxonomic classification as a hierarchical confidence estimation problem rather than a single-rank prediction task. Perseus refines k-mer-level taxonomic signals from Kraken2 using a multi-headed convolutional neural network that estimates calibrated confidence scores for taxonomic correctness at each canonical rank. Using these estimates, Perseus confirms assignments, backs off to higher taxonomic ranks, or abstains when evidence is insufficient, prioritizing correctness and lineage consistency over overly specific assignments. Across simulations of taxonomic novelty and real-world metagenomic datasets, Perseus consistently and substantially reduces the false assignment rate while improving precision and lineage-consistent accuracy. These improvements are most pronounced for long reads and assembled contigs, where spatial context enables reliable discrimination between consistent taxonomic signal and spurious matches.

**Availability and implementation:** Perseus integrates with existing Kraken2 workflows and is available at https://github.com/matnguyen/perseus.

**Supplementary information:** Supplementary data are available online.

## 1 Introduction

Accurate taxonomic classification is a fundamental step in metagenomic studies, enabling characterization of microbial community composition in clinical and environmental samples [25]. However, environmental microbiomes, especially soils, harbor some of the highest microbial diversity on the planet [5], much of which remains uncultivated and taxonomically uncharacterized [20]. Short-read metagenomics has benefited from highly optimized reference databases and k-mer-based classifiers such as the widely used Kraken and Kraken2 algorithms [6, 25], which enable ultrafast taxonomic classification through exact k-mer matching and lowest common ancestor assignment [9, 16]. Kraken2 further reduced memory requirements while supporting substantially larger reference databases, helping standardize k-mer-based classification workflows across diverse metagenomic applications [11, 24].

However, these approaches treat sequences primarily as collections of independent k-mer matches and do not explicitly model how taxonomic evidence is distributed along a sequence. This is particularly problematic for long-read sequencing technologies, which are now seeing widespread adoption in the field. [1, 3, 22, 26]. Several benchmarking studies show that when long reads are used with k-mer-based taxonomic classifiers, even a single mis-assigned region (e.g., a spurious cluster of k-mer matches) can lead to apparently confident but biologically implausible classifications [13, 18]. As a result, long-read datasets often exhibit inflated false positive rates, particularly when the true taxon is absent from the reference database or when related lineages share conserved genomic segments. These errors arise because localized k-mer matches can dominate classification decisions even when the broader sequence context does not support the assigned lineage.

Several strategies have been proposed to address this problem, including expanded reference databases [6, 15], confidence thresholds [25], and post-processing filters [14]. However, simply enlarging reference databases is increasingly difficult: many environmental lineages lack high-quality representatives [17]. Confidence-based heuristics require dataset-specific, hand-tuned thresholds that vary across sequencing platforms and environments, often trading sensitivity for specificity without understanding why a prediction should be rejected [10]. More broadly, current methods struggle to exploit the rich spatial and hierarchical signal present within a sequence and typically treat each sequence as an unstructured “bag of votes”. Crucially, they generally ignore the hierarchical nature of taxonomy: a misclassification at the species level may still be partially correct at the genus or family level. However, most filters make binary decisions at a single rank without leveraging lineage-consistent evidence across the taxonomic hierarchy.

Here, we present **Perseus**, a lineage-aware convolutional neural network that refines Kraken2 classifications for long reads and contigs by learning spatially structured patterns of k-mer assignments along each sequence. For each sequence-taxon pair obtained from Kraken2, Perseus extracts a 22-channel representation that captures raw k-mer signals, in-lineage and out-of-lineage evidence, and descendant/ancestor contributions. The model then estimates confidence probabilities at seven canonical ranks simultaneously, allowing it to model the taxonomic hierarchy and identify the deepest rank supported by evidence. From a machine learning perspective, Perseus can be viewed as a hierarchical confidence estimation model operating over taxonomic label space, reframing taxonomic classification as estimating the deepest lineage supported by evidence rather than predicting a single fixed rank.

Conceptually, Perseus separates evidence accumulation from confidence estimation in taxonomic classification: Kraken2 aggregates k-mer matches, while Perseus evaluates whether those matches form a coherent lineage-consistent pattern of evidence. Rather than treating k-mers as independent votes, Perseus performs lineage-aware confidence estimation using spatial and hierarchical consistency of the k-mer evidence, modeling how taxonomic evidence is distributed along each sequence. This allows Perseus to identify the deepest taxonomic rank supported by structured evidence, backing off or abstaining when fine-rank assignments are driven primarily by sparse or localized matches.

## 2 Methods

### 2.1 Training and Evaluation Data

We construct four experimental datasets derived from Kraken2’s standard bacterial reference database to systematically evaluate classification robustness under varying degrees of taxonomic novelty. Three datasets are designed to model increasing evolutionary distance between query sequences and the reference database by excluding taxa at the species, genus, and family level, respectively. To reduce bias from fragmented or low-quality references, we first filter the bacterial database to retain only genomes longer than 1 Mbp along with their associated plasmids. Then, for each exclusion level, we stratify taxa by phylum and remove 250 species, 150 genera, or 100 families, ensuring that excluded taxa span broad phylogenetic diversity rather than being concentrated within a small number of clades. In addition to these exclusion datasets, we construct a core inclusion dataset consisting of species that were retained across all three exclusion datasets. Each of the four datasets is then built as an independent Kraken2 reference database.

We then use MeSS [4] to generate simulated long reads for model training and evaluation. For each dataset, both included and excluded taxa are partitioned into three non-overlapping sets for training, validation, and testing, and independent simulation runs are generated for each split. To capture a range of data characteristics, we simulate four sequencing conditions per dataset: (i) high-quality PacBio HiFi reads (8-22 kbp read lengths), (ii) shorter and lower-quality PacBio HiFi reads (1-7 kbp read lengths), (iii) Oxford Nanopore reads with lengths ranging from 2–100 kbp, and (iv) a long-tailed Oxford Nanopore distribution with reads up to 200 kbp. During training, the model is trained jointly on data from all four sequencing conditions, while validation and testing are performed on held-out simulation runs. To reflect realistic abundance heterogeneity in metagenomic samples, simulated coverage ranges from 0.5× to 300×. Using these data, the model is trained once on simulated datasets and applied without retraining to all evaluation datasets.

In addition to these synthetic datasets, we also evaluate Perseus’ performance on the plant-associated and marine CAMI II datasets. Furthermore, we also evaluate the model on real data. We test on a PacBio HiFi run of the ZymoBIOMICS Gut Microbiome Standard D6331 (accession: SRR13128014) to assess performance on a real long-read dataset with known species composition. Finally, we apply Perseus to an Oxford Nanopore soil metagenome from the Schönbuch Forest [2] as an exploratory case study of performance on complex environmental microbiomes.

### 2.2 Feature Encoding

Kraken2 is a k-mer-based taxonomic classifier that assigns sequences based on k-mer matches to a reference database. It uses a compact hash table of minimizers and maps each k-mer in a sequence to the lowest common ancestor of all reference genomes containing that k-mer. A sequence’s final classification is determined by aggregating the taxonomic labels of its k-mers and selecting the taxon with the highest count. The primary output of Kraken2 is a tab-delimited file with one row per sequence and five columns: (1) the classification status (C for classified and U for unclassified), (2) the sequence/read name, (3) the assigned taxonomic ID, (4) the sequence length, and (5) a k-mer report string encoding the number of consecutive k-mers assigned to a taxon.

Perseus reframes taxonomic filtering as a lineage-aware confidence estimation problem. Rather than treating k-mers independently, we model spatial consistency of k-mer assignments along each sequence. We encode features for the Perseus model by post-processing the Kraken2 output (Figure 1). For each sequence *s*, Kraken2 produces a list of k-mer-level taxonomic assignments for each position *p*: *κ*_*s*_(*p*) ∈ 𝒯 where 𝒯 is the set of NCBI taxonomic IDs, and *κ*_*s*_(*p*) = 0 for unclassified k-mers and *κ*_*s*_(*p*) = *A* for ambiguous k-mers. We begin by dividing the k-mer report string for each sequence into *T* non-overlapping bins of length Δ = 1000bp, so that bin *B*_*t*_ covers positions [(*t* − 1)Δ, *t*Δ) for *t* = 1, …, *T*. For each taxon *c* that appears in the Kraken2 output for a sequence, we form a sequence-taxon pair (*s, c*). For each pair, we compute *C* = 22 lineage-aware feature channels over the *T* bins, resulting in a structured representation of the spatial distribution of k-mer evidence for taxon *c* along the sequence.

**Figure 1.**
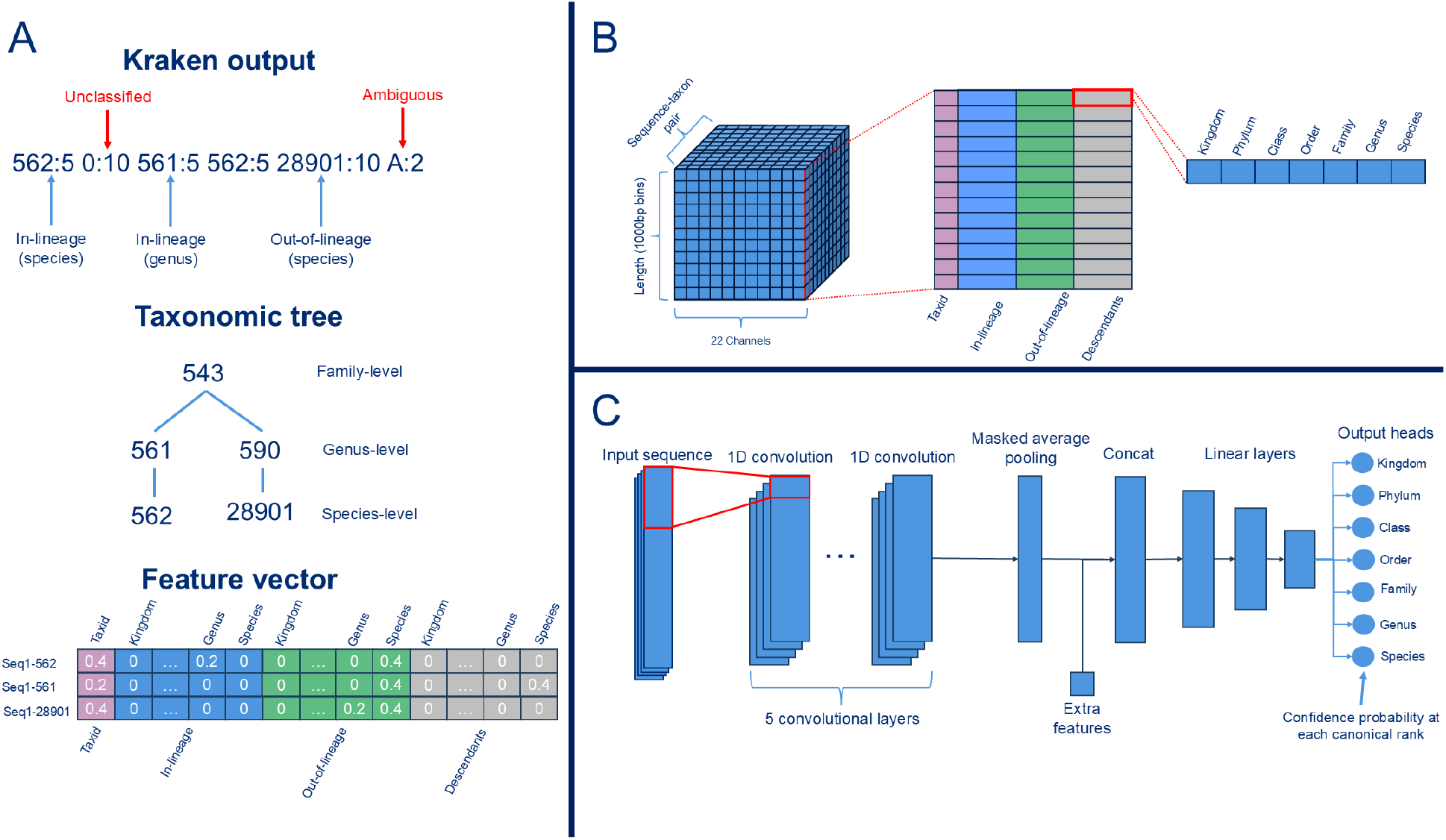
Feature encoding and model architecture of Perseus. (A) Kraken2 k-mer assignments are aggregated into 1 kb bins along each sequence to form a multi-channel feature tensor. For each sequence–taxon pair, channels encode the distribution of k-mers assigned to the candidate taxon (taxid *c*), its ancestors (in-lineage), its descendants, and out-of-lineage taxa across the canonical ranks (kingdom → species). This representation preserves the spatial organization of taxonomic signal along the sequence. (B) Example of encoding Kraken2 output into the feature tensor. We illustrate the k-mer assignments from a sequence Kraken2 initially assigned as taxon 562. For each sequence-taxon pair, Perseus constructs feature vectors that capture lineage-consistent and lineage-inconsistent evidence. In this example, the mapping for each k-mer includes taxonomic ids 562 (*Escherichia coli*), 561 (*Escherichia*), and 28901 (*Salmonella enterica*). For the first vector (Seq1-562), 543 lies in-lineage at the genus level, while 28901 is out-of-lineage at the species level. For the second vector (Seq1-561), 562 is a descendant at the species level, and 28901 remains out-of-lineage at the species level. Finally, for the third vector (Seq1-28901), 562 is out-of-lineage at the species level, and 561 is out-of-lineage at the genus level. Unclassified and ambiguous k-mers are discareded during feature construction. (C) The feature tensor is processed by a stack of 1D convolutional layers to learn local spatial patterns of taxonomic consistency. Masked average pooling produces a fixed-length embedding for variable-length inputs, which is concatenated with extra features (e.g., sequence length) and passed through fully connected layers. The network terminates in multiple output heads, each predicting a confidence probability for the correctness of the assignment at a specific canonical taxonomic rank (kingdom through species).

Each feature channel represents the proportion of k-mer assignments within each bin. For a sequence *s*, we define the number of assigned k-mers in bin *B*_*t*_ as

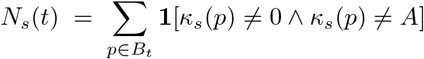

where we discard any k-mers whose Kraken2 assignment is unclassified (*κ*_*s*_(*p*) = 0) or ambiguous (*κ*_*s*_(*p*) = *A*). All feature values are set to zero whenever *N*_*s*_(*t*) = 0.

To capture both spatial and hierarchical consistency of taxonomic evidence, we group feature channels into four categories according to their taxonomic relationship to candidate taxon *c*: raw evidence, in-lineage evidence, out-of-lineage evidence, and descendant evidence. The raw evidence channel measures the proportion of k-mers in bin *B*_*t*_ that are assigned exactly to taxon *c*:

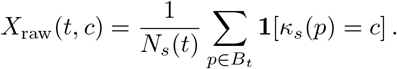

In-lineage evidence captures k-mers in the bin that Kraken2 assigns to any ancestor of *c*, reflecting higher-level rank support for the same lineage. Let anc(*c*) denote the set of all ancestor taxa of *c* such that anc(*c*) ∈ {kingdom, phylum, class, order, family, genus, species}. Then

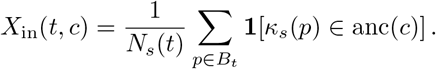

Out-of-lineage evidence represents the proportion of k-mers assigned to neither *c* nor to any ancestor of *c*, which quantifies conflicting signals:

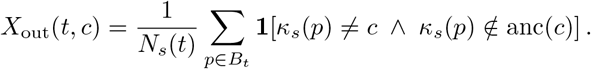

Finally, descendant evidence measures support for more specific taxa below *c*. Let desc(*c*) denote all strict descendants of *c* in the taxonomic tree. Then

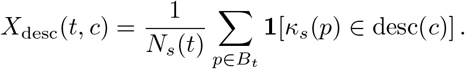

Together, these channels capture the distribution of lineage-consistent and lineage-inconsistent signals across the sequence to form a structure featured tensor representation *X*_*s,c*_ ∈ ℝ^*C*×*T*^ used by the convolutional neural network.

### 2.3 Model Architecture

To capture spatial patterns of taxonomic evidence while modeling the hierarchical structure of taxonomic labels, we use a multi-headed one-dimensional convolutional neural network that operates on the feature tensor *X*_*s,c*_ ∈ ℝ^*C*×*T*^ for each sequence-taxon pair. The network begins with five 1D convolutional blocks that extract local spatial patterns across adjacent bins. We then apply masked average pooling to obtain a fixed-size representation from the convolutional outputs while ignoring padded positions to support variable-length inputs. We then concatenate the sequence length with the pooled vector to form a single embedding. This joint embedding is processed through three fully connected layers, after which the final layer branches into seven independent output heads corresponding to the canonical taxonomic ranks from kingdom through species. During training, we apply focal loss independently to each head to emphasize difficult, low-confidence examples and mitigate class imbalance across ranks, especially at the highest taxonomic ranks where Kraken2 is already precise. For sequences lacking labels at certain ranks, loss contributions from those heads are masked, and the total loss is normalized by the number of supervised rank predictions in each batch.

Because the raw sigmoid outputs for each head are often poorly calibrated, we apply post-hoc isotonic regression to convert CNN scores into reliable probability estimates. A validation dataset consisting of 20% of the training data, stratified by species, was used to collect pairs of raw CNN scores and binary correctness labels for each canonical rank. For each rank, we fit an independent isotonic regression model, producing a monotone mapping from raw scores to empirical correctness frequencies. This procedure places outputs on a calibrated probability scale, such that each head produces a single probability *y*_(*s,c*)_(*r*) ∈ [0, 1], representing the confidence that taxon *c* is correct at rank *r*.

### 2.4 Performance Metrics

We measure the accuracy of Kraken before and after apply Perseus at a per-read level. Because reads generated by MeSS can be traced back to their corresponding reference genome, the ground-truth taxonomic label is known for each simulated read. Most of our performance metrics are derived from the definitions used in Kraken benchmarking studies [25], and consist of true positives (TP), lineage-correct positives (LCP), false positives (FP), and false negatives (FN). For a ground-truth species, a TP occurs when the classification matches the ground truth. A LCP occurs when the classification is an ancestor of the true species in the taxonomic lineage. A FP is an incorrect classification that neither matches the true species, nor lies within its lineage. An FN occurs when Kraken2 or Perseus fails to assign any classification. All sequences in our inclusion/exclusion datasets should match the reference database at least at the phylum level, and thus true negatives are not defined in this evaluation setting.

Using these categories, we define the call rate, as the proportion of sequences receiving a taxonomic assignment at any rank: (TP + LCP + FP) / (TP + LCP + FP + FN). We define the positive predictive value (PPV) as the proportion of classifications that were TP (excluding lineage-correct assignments): TP / (TP + FP). We also define lineage-correct PPV, which accounts for lineage-correct assignments: (TP + LCP) / (TP + LCP + FP). We define the false assignment rate (FAR) as the proportion of assignments that are incorrect: FP / (FP + TP + LCP). Recall is defined as the proportion of correctly called sequences: TP / (TP + LCP + FP + FN). The F1-score is defined as the harmonic mean of the recall and PPV. Given that the F1-score does not take into account the lineage-correctness of assignments, we also define the lineage-correct recall the proportion of sequences that were either TP or LCP: (TP + LCP) / (TP + LCP + FP + FN). Relatedly, the lineage-correct F1-score (LC-F1) is defined as the harmonic mean of the LC-recall and LC-PPV.

## 3 Results

### 3.1 Taxonomic Rank Removal Experiment

We performed a taxonomic rank removal experiment to assess failure modes of Kraken2 when classifying sequences from novel genomes. We first constructed the standard Kraken2 bacterial database, which contains all RefSeq complete bacterial genomes. We then queried the *E. coli* O104:H4 substrain. Next, we removed all sequences of the *E. coli* species, re-built the Kraken2 database, and re-queried the O104:H4 substrain. We then repeated this procedure to remove all sequences of the genus *Escherichia* and again after removing all sequences from the family Enterobacteriaceae.

At each level, we count the number of k-mers that fall within the following categories: correct, for k-mers that directly match to the O104:H4 substrain; lineage-correct, for k-mers classified at a higher taxonomic rank within the lineage of the substrain; incorrect, for k-mers that are not lineage-consistent; and unclassified, for k-mers that are unclassified or ambiguous.

As shown in Figure 2, the majority of k-mers map no more specifically than the family level, and just 0.02% of k-mers are assigned to the correct strain. Consequently, baseline strain-level calls are typically supported by only a small number of k-mers. This reflects the high sequence similarity shared among closely related species and the limited availability of truly strain-specific k-mers. As a result, fine-grained predictions are based on sparse evidence and become particularly error-prone when exact matches are absent from the database.

**Figure 2.**
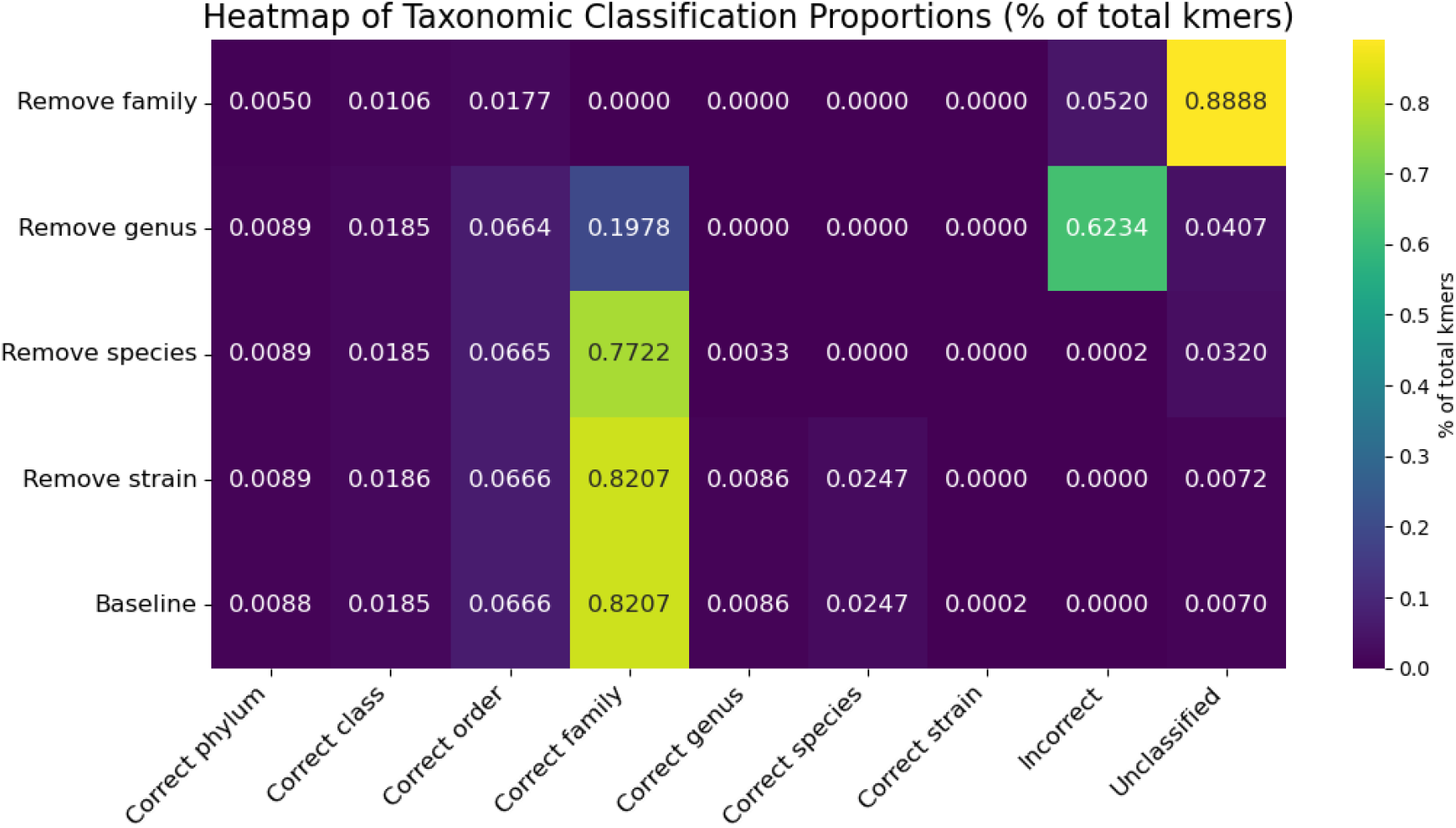
Distribution of k-mer–level taxonomic assignments under systematic rank removal. Heatmap showing the proportion of k-mers assigned to each classification category across baseline and rank-removal Kraken2 databases. Kraken2 outputs a single taxonomic match for each k-mer. Rows correspond to the taxonomic rank removed from the reference database, and columns indicate the finest taxonomic level to which each k-mer was assigned. Correct (accession) denotes k-mers assigned to the true O104:H4 strain; Correct *i* denotes assignments to rank *i* within the true lineage (domain → species); Incorrect denotes off-lineage assignments; and Unclassified denotes k-mers receiving no assignment (or ambiguous k-mers). Values represent the fraction of total k-mers. For example, any k-mers assigned to Enterobacteriaceae would fall into the “Correct genus” category. The results illustrate how removing lower taxonomic ranks shifts k-mer support toward higher lineage-consistent ranks and increases unclassified or incorrect assignments.

This behavior becomes evident when we remove the entire *Escherichia* genus: the misclassification rate increases to 62%. When we remove the entire Enterobacteriaceae family, the vast majority of k-mers become unclassified (89%), representing a desirable outcome for genuinely novel sequences. However, even in this idealized condition, Kraken2 still produces a 5% misclassification rate, which is higher than when only the species or genus are removed. Under the family-exclusion scenario, we observe that misclassified k-mers are not randomly distributed across the taxonomy. Instead, 9.7% of incorrect k-mers mapped broadly to the Morganellaceae family. At the species-level, the frequent false positive assignment was *Proteus mirabilis* accounting for 6.1% of misclassified k-mers. This demonstrates that even when most k-mers become unclassified, the remaining false positives may match to neighboring but incorrect taxa, highlighting the susceptibility of exact k-mer-based classifiers under database incompleteness. Together, these results demonstrate that a substantial fraction of Kraken2’s taxonomic assignments can arise from false positive evidence, driven by spuriously mapped k-mers, especially under incomplete or underspecified reference databases.

### 3.2 Source of Misclassification

To better understand the source of misclassifications, we first align the O104:H4 substrain of *E. coli* against the *Proteus mirabilis* HI4320 strain reference genome using Minimap2 [8]. We identified 13 local alignments (median length 1147 bp). Annotation of the aligned regions in O104:H4 showed that they corresponded to rRNA operons. No conserved protein-coding regions were observed outside of these ribosomal loci, indicating that the *Proteus* signal is driven by highly conserved ribosomal sequences rather than broader genomic similarity.

To measure the extent of sequence conservation among bacterial genomes, we randomly sampled 100 bacterial reference genomes from RefSeq, and performed all pairwise whole-genome alignments using MUMmer4 [12]. This analysis extends the previous case study by examining sequence conservation across a diverse set of bacterial genomes, providing a broader context for the locus-specific similarity observed between O104:H4 and *Proteus mirabilis*.

Across all 9000 genome pairs, we observe 73% of pairs (6604 pairs) with at least one conserved segment of length ≥ 50 bp, which implies the presence of multiple adjacent k-mer matches (at least 50-31+1=20 for 31-mers) in Kraken2. Moreover, 24% of pairs (2154 pairs) shared at least one conserved segment ≥ 200 bp. The mean number of conserved regions per genome pair was 170, while the median was 4, indicating that most genome pairs only share limited number of conserved regions ≥ 50 bp while a small subset of genome pairs contain up to 12,000 conserved regions. These conserved regions were typically short: conserved regions ≥ 50 bp have a mean length of 256 bp, and a median length of 73 bp. This shows the typical conserved region just exceeds the default k-mer length used by classifiers like Kraken2 (31 bp), meaning they span multiple adjacent k-mers and providing a mechanistic explanation for how short conserved regions can generate misleading k-mer evidence. In contrast, conserved regions ≥ 200 bp have a mean length of 3.03 kb and a median length of 1.28 kb.

To consistently annotate the aligned regions, each genome was independently annotated using Bakta [21], to obtain GFF3 feature annotations including protein-coding genes, structural RNA elements, and other genomic features. Aligned regions were intersected with genome annotations using bedtools intersect [19]. For alignment intervals overlapping multiple genomic features, the feature with the largest overlap was selected to assign a single annotation per aligned region. Aligned regions were classified into broad categories (protein-coding, RNA, mobile genetic element, or intergenic) based on feature type.

To assess the relationship between alignment length and functional conservation, aligned regions were stratified by alignment length into discrete bins between 50 bp and 200 bp, and the fraction overlapping annotated conservation features was computed for each bin. Summary statistics were calculated across all genome pairs, enabling systematic assessment of conserved regions capable of generating k-mer matches that may lead to spurious taxonomic assignments.

Among conserved regions ≥ 200 bp, 93% overlap with annotated features. Of these annotated regions (184,916), 143,206 (77.44%) corresponded to protein-coding regions, 27,348 (14.79%) to structured RNA elements, and 14362 (7.77%) to mobile genetic elements. Among protein-coding genes, conserved regions were enriched for essential housekeeping genes such as those for transcription and translation machinery (e.g. RNA polymerase subunits and elongation factors), metabolic enzymes (e.g. nitrate reductase and transketolase), and bacterial molecular chaperones (e.g. GroEL, DnaK). Interestingly, the most highly enriched annotation of protein-coding regions was for hypothetical proteins. Structured RNA elements were also common, with long conserved alignments overlapping with rRNA operons, with 23S rRNAs being the most enriched, followed by 16S. Multiple tRNA species and the transfer-messenger RNA SsrA were also prominent. Finally, a large fraction of long conserved regions correspond to mobile genetic elements, predominantly insertion sequences (IS) family transposases, with the most common being the IS1 family.

These results indicate that taxonomic misclassification often arises from short, spatially localized conserved regions that generate clusters of k-mer matches, motivating approaches that model the spatial distribution and lineage consistency of k-mer evidence along sequences.

### 3.3 Effect of Confidence Score Thresholding

In Kraken2, the confidence parameter (`--confidence`) specifies the minimum fraction of k-mer evidence that must support a taxon for that taxon to be reported, enforcing a minimum threshold for assignment strength and causing low-support sequences to be reassigned to higher ranks or marked as unclassified. To evaluate whether tuning Kraken2’s internal confidence parameter mitigates over-specific assignments, we measured classification performance across a range of confidence thresholds (0.0 to 1.0 in increments of 0.01). For each confidence value, reads were classified independently and evaluated using the same experimental inclusion/exclusion datasets (Supplementary Figure S1).

As expected, increasing the confidence threshold monotonically reduces the overall call rate across all datasets. In the inclusion dataset, the call rate remained high across a broad range of thresholds but dropped sharply at extreme values (*>* 0.9). In contrast, exclusion datasets exhibited lower baseline call rates and declined more rapidly with increasing threshold, especially under family exclusion. PPV improves substantially when moving from a confidence of 0.0 to low non-zero thresholds, but changes only modestly across intermediate thresholds. At very high confidence values, PPV fluctuates due to a large reduction in classified reads. Similar trends are observed for LC-PPV, which increases gradually with increasing confidence scores and approaches maximal values only at thresholds where large numbers of reads become unclassified. The FAR decreases as the confidence threshold increases, but plateaus at intermediate thresholds. At extreme thresholds, FAR approaches zero because most reads are assignments to higher taxonomic ranks or completely unclassified. Consistent with these trends, F1 score declined steadily as confidence increases, showing the tradeoff between precision gains and reduced recall.

These results show that while increasing the Kraken2 confidence threshold reduces false assignments, it does so primarily by aggressively reducing call rate. Across a broad intermediate range of thresholds, exclusion datasets retain substantial false assignment rates, indicating that confidence filtering alone does not eliminate over-specific classifications supported by localized conserved k-mer evidence. Therefore, we conclude that the Kraken confidence threshold enforces a blunt precision-recall tradeoff without accounting for the spatial or hierarchical consistency of k-mer evidence.

### 3.4 Evaluation on Inclusion/Exclusion Data

We next evaluate Perseus on both simulated and benchmark metagenomic datasets, including MeSS-simulated reads from our core inclusion dataset and three exclusion datasets, as well as simulated reads and assembled contigs from the CAMI II marine and plant-associated benchmarks. The inclusion and exclusion datasets allow us to characterize classifier behavior under increasing degrees of reference database incompleteness, corresponding to family-, genus-, and species-level exclusions. In contrast, the CAMI II datasets enable evaluation of Perseus on both read-level and contig-level metagenomic data. For each benchmark, we compare baseline Kraken2 classifications with those obtained after Perseus post-processing (Supplementary Table S1).

We evaluate the performance of Kraken2, Kraken2 post-filtered with Perseus, Centrifuger [23] and Metabuli [7] in Figure 3. Both Centrifuger and Metabuli are reference-based classification methods: Centrifuger employs an FM-index for exact nucleotide sequence matching while Metabuli uses a hybrid k-mer framework (“metamers”) that leverages amino acid conservation to improve sensitivity to homologous sequences despite nucleotide variation. Across all inclusion/exclusion datasets, Kraken2 maintains a consistently high call rate but exhibits elevated FAR, especially under higher-rank exclusion scenarios. In the family exclusion dataset, FAR approaches 50%, highlighting the susceptibility to reference incompleteness. In contrast, Perseus substantially increases LC-PPV (Panel A) and reduces FAR (Panel B) especially in exclusion experiments. These gains reflect more conservative, lineage-consistent predictions. While call rate decreases after filtering, this reduction reflects deliberate abstention on uncertain fine-rank assignments, rather than erroneous classifications.

**Figure 3.**
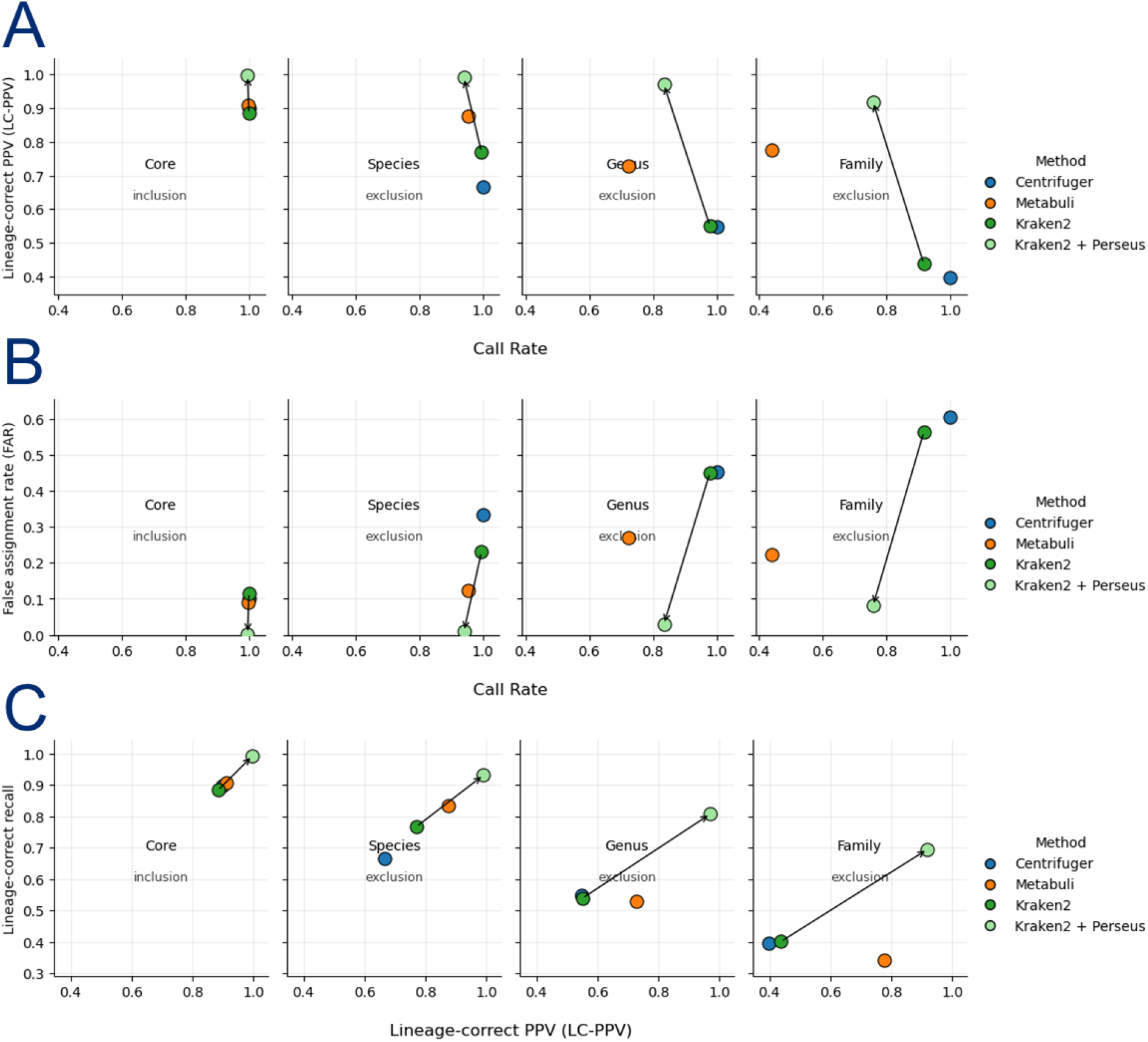
Performance of Perseus across inclusion/exclusion simulations and CAMI benchmarks. Performance on inclusion/exclusion datasets. Each column corresponds to an experiment (core inclusion, species exclusion, genus exclusion, family exclusion) and points denote methods (Centrifuger, Metabuli, Kraken2 and Kraken2 filtered with Perseus). Arrows connect Kraken2 to Perseus-filtered points to highlight the effect of post-processing. (A) Lineage-correct positive predictive value (LC-PPV) versus call rate. Perseus consistently increases LC-PPV relative to Kraken2 across all datasets, with the largest gains observed under higher-rank exclusions, while maintaining comparable call rates. (B) False assignment rate (FAR) versus call rate. Perseus substantially reduces FAR compared to Kraken2, particularly in species and genus exclusion datasets, demonstrating robustness to missing references. (C) Lineage-correct recall versus LC-PPV. Perseus shifts performance toward higher precision-recall tradeoffs, especially in genus and family exclusion datasets. This reflects Perseus’ more conservative but lineage-consistent predictions.

Panel C shows the precision-recall tradeoff of Perseus, demonstrating that it shifts predictions towards higher LC-PPV while maintaining competitive lineage-correct recall. This behavior demonstrates that Perseus backs off to higher taxonomic ranks within the correct lineage rather than making overconfident species-level assignments. Importantly, Kraken2 post-filtered with Perseus consistently outperforms Centrifuger and Metabuli in LC-PPV, FAR, and LC-F1 across nearly all experiments. In higher-rank exclusion scenarios, the Perseus filtered Kraken2 pipeline achieves the strongest precision-recall tradeoff. Figure 4 (Panel A) shows that LC-F1 is consistently highest after Perseus post-processing, indicating improved robustness in novel-genome scenarios.

**Figure 4.**
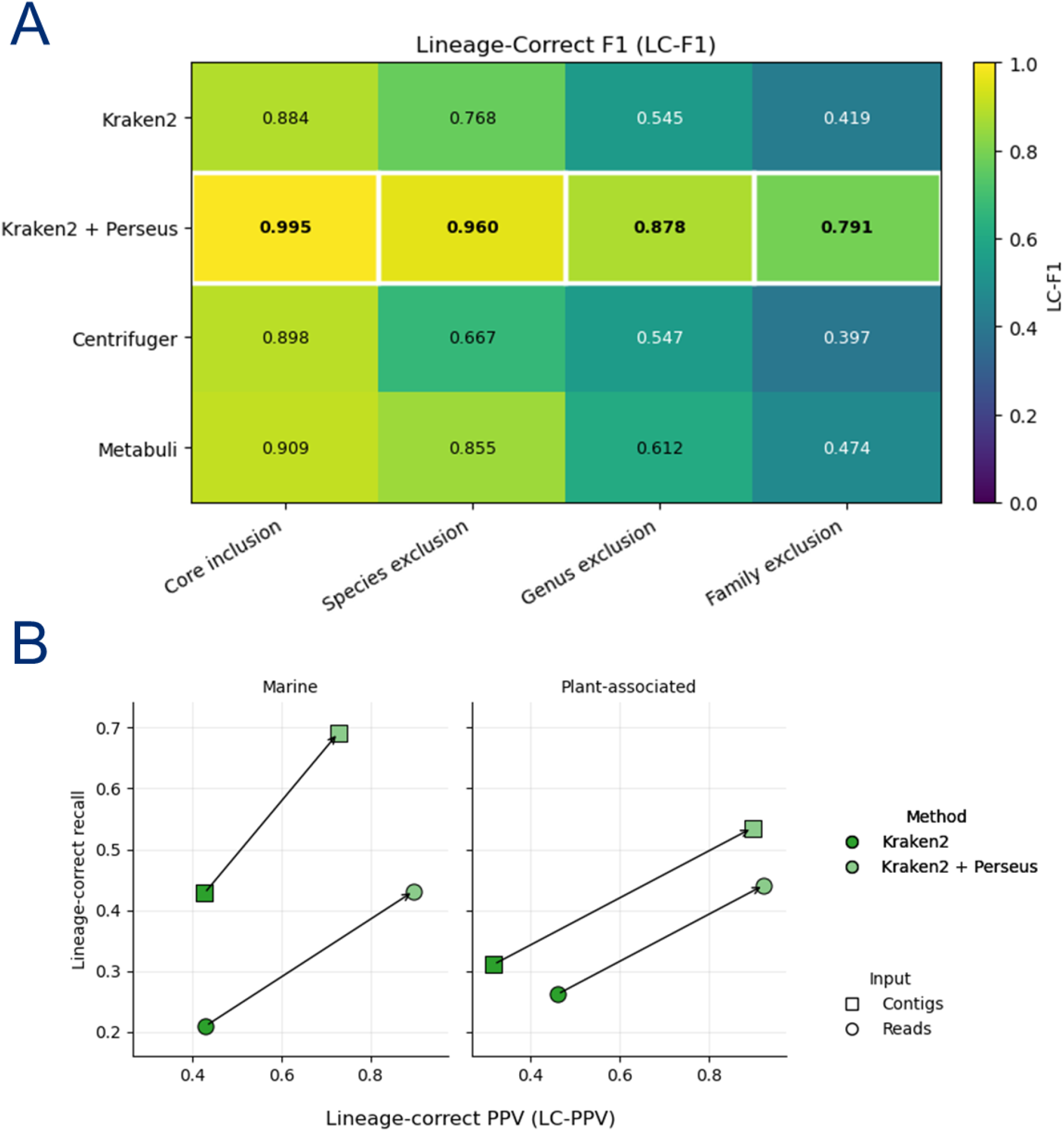
Performance of Perseus across inclusion/exclusion simulations and CAMI benchmarks. (A) Heatmap of lineage-correct F1 (LC-F1) across inclusion/exclusion datasets. LC-F1 is consistently highest after post-processing with Perseus, with the greatest improvements under species, genus, and family exclusion datasets, demonstrating improved performance in novel-genome scenarios. (B) Results on CAMI II marine and plant-associated datasets (reads and contigs). Points show lineage-correct recall versus LC-PPV. Perseus improves LC-PPV and lineage-correct recall across both datasets and input types, with stronger gains observed on contigs.

As seen in Figure 5, Perseus suppresses low-confidence, spurious fine-rank assignments by converting many FPs into lineage-consistent higher rank calls (now LCP) or abstentions (FN/unclassified), while retaining a large portion of TPs. We also see a large portion of FPs being reclassified as LCP, indicating that many apparent FPs occur because of over-specific assignments, rather than completely incorrect high-level assignments. These results show Perseus’ conservative behavior, prioritizing correctness and lineage-consistency, especially at higher taxonomic levels, rather than making overconfident fine-rank assignments. Importantly, Perseus does not simply reduce call rate uniformly; instead, many false-positive species-level assignments are converted into lineage-consistent higher-rank calls rather than being discarded entirely, demonstrating structured, rank-aware refinement rather than indiscriminate abstention.

**Figure 5.**
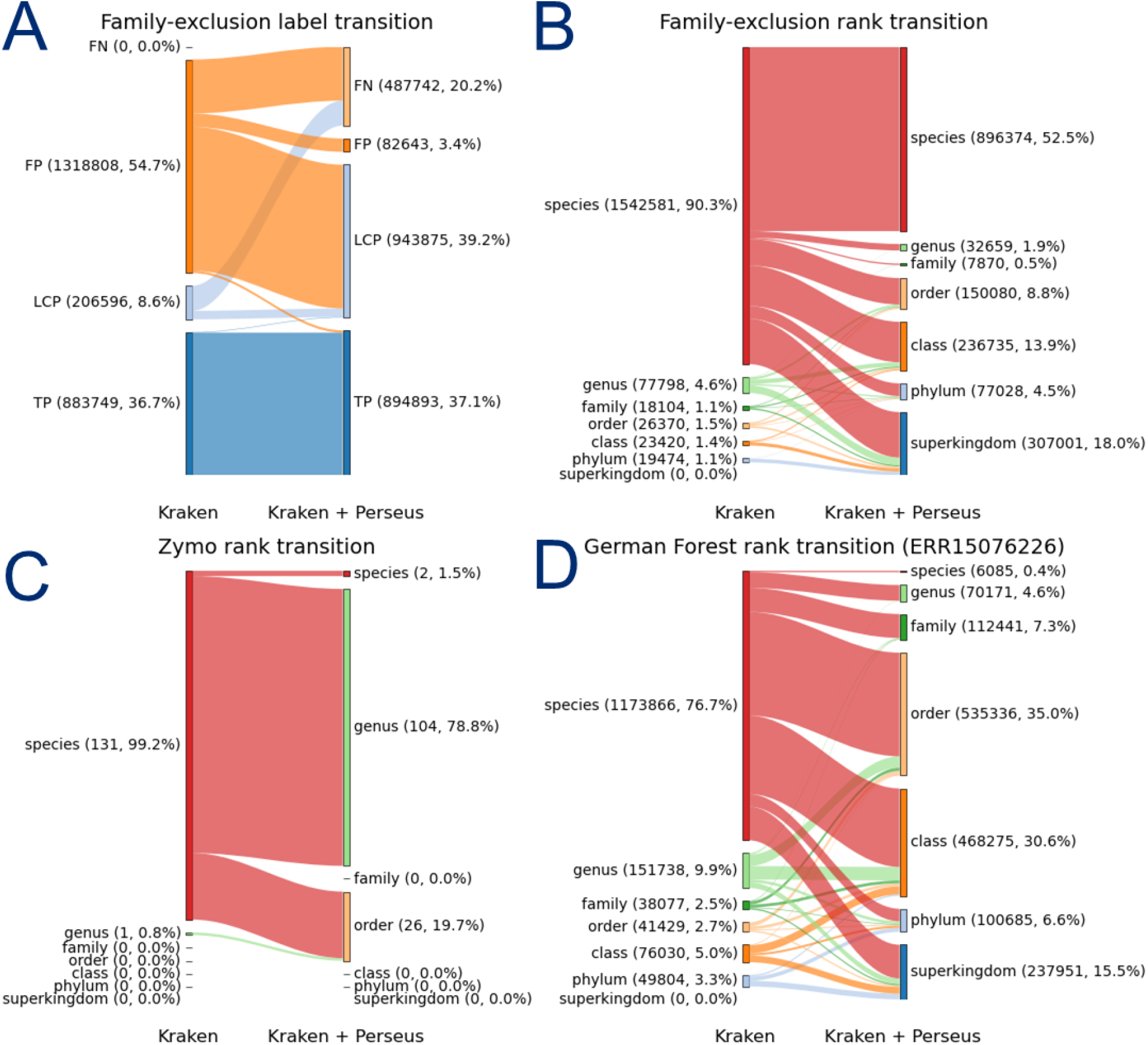
Redistribution of taxonomic assignments before and after Perseus filtering. (A) In the family-exclusion experiment, Kraken2 predictions are categorized as TP, LCP, and FP. Perseus substantially reduces FPs by converting many low-confidence spurious fine-rank errors into higher-rank lineage-consistent calls or abstentions, while retaining the majority of TPs. We also see a large portion of FPs being reclassified as LCP, indicating that many apparent FPs occur because of over-specific assignments, rather than completely incorrect high-level assignments. (B) Rank-level redistribution in the family-exclusion setting shows back-off from species to order, class, phylum, and superkingdom, reflecting suppression of unsupported fine-grained assignments. (C) On the Zymo dataset, Perseus shifts a large fraction of species-level predictions to higher ranks, reducing over-specific calls and increasing conservative, lineage-consistent assignments. (D) Similar trends are observed for the German Schönbuch Forest microbiome (ERR15076226), where many species-level predictions are reassigned to order or class, illustrating rank-aware filtering that favors lineage consistency over overconfident specificity.

### 3.5 Evaluation on CAMI Datasets

On the CAMI II benchmark datasets, we see universal improvements in the PPV, LC-PPV, and FAR after filtering with Perseus. For the marine dataset, call rate modestly decreases for both reads and contigs. In contrast, for the plant-associated reads, Perseus nearly eliminates all false assignments (very low FAR), but aggressively abstains, resulting in a low call rate and F1-score. This is likely because the long-reads from the dataset are extremely short, with a mean length of only 2kb. Notably, when applied to the plant-associated contigs, Perseus is much less conservative, with improvements in PPV, LC-PPV, and FAR, while maintaining a more balanced call rate. These results indicate two key properties: first, Perseus is more effective for longer reads, or even contigs, when more spatial k-mer context is available. Secondly, in highly novel or under-represented environments, Perseus will prioritize conservative predictions, and abstain rather than overconfidently misclassifying.

On the CAMI II benchmark datasets (Panel B of Figure 4), Perseus consistently shifts performance toward improved lineage-aware precision and recall relative to Kraken2. In the marine dataset, both reads and contigs show clear gains in LC-PPV and lineage-correct recall after filtering. This improvement is particularly pronounced for contigs, where LC-PPV increases substantially and recall improves from 0.43 to 0.69. This shows Perseus is able to leverage longer sequence context to refine taxonomic assignments without sacrificing sensitivity. In the plant-associated data, improvements are again observed for both reads and contigs. For reads, Perseus increases LC-PPV while also improving lineage-correct recall. For contigs, the gains are even stronger, with both LC-PPV and recall increasing simultaneously showing a favorable shift in precision-recall.

### 3.6 Evaluation on Real-world Data

We also applied Perseus to PacBio HiFi reads generated from the ZymoBIOMICS Gut Microbiome Standard (SRR13128014) (Panel C in Figure 5). On this mock community, Kraken2 already achieves strong species-level performance because all 21 species present are well represented in reference databases and are generally quite distinct from each other. For the subset of TP assignments, Perseus backs off from species- or strain-level assignments to the higher ranks, most commonly at the genus level. This is because there is little k-mer evidence that strongly supports lineage consistency and spatial support at the species-level. Therefore, Perseus favors higher-rank assignments that are robustly supported, even if fine-rank predictions are potentially correct. Consequently Perseus effectively suppresses Kraken2 errors: Perseus filters out most of the incorrect species-level classification from Kraken2, by backing-off to the correct higher-level ranks. Together, these results highlight that Perseus trades taxonomic resolution for improved reliability at higher ranks such as at the genus or family level.

We next applied Perseus to the individual Oxford Nanopore reads from a German Schönbuch Forest soil sample (Panel D in Figure 5). We also used this sample for runtime and memory benchmarks (Supplementary Table S2). These data are highly diverse and contain uneven coverage, making for a particularly challenging problem for k-mer-based taxonomic classification. In baseline Kraken2 results, species-level assignments are the most common among reads, with over 1,100,000 reads receiving species-level taxonomic classifications. However, after Perseus filtering, only 5974 reads retained species-level assignments, which is more than a 2 orders of magnitude reduction. Instead of discarding these reads as unclassified, Perseus conservatively assigns most of them to higher taxonomic ranks, especially at the order (35%) or class (30%) levels, with few high rank assignments remaining.

To assess whether the reduction in species-level assignment corresponded to improved specificity rather than loss of signal, we examined the k-mer support for the retained classifications. In baseline Kraken2 results, species-level assignments were frequently supported only by a small number of k-mers (median = 5, mean = 14), despite the reads typically exceeding kilobases in length. After filtering with Perseus, the remaining species-level assignments have much stronger support, with a median of 187 and a mean of 389 k-mers matching the classified taxon. We also continue to see this trend after normalizing for read length, with length-normalized k-mer support increasing substantially after filtering with Perseus.

To further investigate the biological validity of Perseus’ back-off behavior, we evaluated alignment support using Minimap2 [8] for different groups of classifications in the sample. For reads originally assigned at the species level by Kraken2, we grouped predictions by the final rank selected by Perseus, then computed the alignment rate to the assigned reference genome and the median query coverage among aligned reads (Supplementary Figure S2). Reads retained at the species or genus level have nearly complete alignment rates and high median query coverage. This indicates a strong concordance with the matched reference genomes. However, reads that Perseus assigns to higher taxonomic ranks (i.e. family and above) show substantially lower alignment rates and drastically lower coverage. This monotonic decline suggests that Perseus preserves fine-rank assignments only when there is robust sequence evidence, while backing off in cases where reference-level support is weak or incomplete. These results provide an alignment-based validation to Perseus’ regime, and demonstrates that Perseus’ confidence modeling yields biologically consistent taxonomic refinement on real-world microbiome data by preserving well-supported assignments while systematically suppressing spurious fine-rank predictions.

## 4 Discussion

Accurate taxonomic classification of long-read metagenomic data remains challenging, particularly for complex environmental microbiomes where many organisms lack close representatives in reference databases. While k-mer-based classifiers such as Kraken2 enable extremely fast taxonomic assignment, they frequently produce overly specific predictions when applied to long reads or assembled contigs. Our analyses show that these errors commonly arise from sparse clusters of conserved k-mers originating from shared genomic elements such as housekeeping genes, rRNA loci, or mobile genetic elements. These localized signals can spuriously support fine-rank assignments even when the broader sequence context does not support the inferred lineage. Furthermore, our results highlight that these errors cannot be effectively addressed by Kraken2’s internal confidence threshold, which reduces false assignments primarily by bluntly increasing abstention rather than determining whether the evidence supports a deeper lineage.

To address this limitation, we introduced Perseus, a lineage-aware post-processing framework that refines Kraken2 classifications by modeling the spatial distribution and hierarchical consistency of k-mer evidence along each sequence. Across simulated inclusion/exclusion experiments, CAMI benchmarks, and real-world datasets, Perseus consistently reduced false positive assignments while preserving lineage-consistent higher-rank classifications. These improvements were especially pronounced in scenarios with high rates of taxonomic novelty, where reference incompleteness increases the risk of over-specific assignments. Importantly, many false-positive species-level assignments are converted into lineage-consistent higher-rank predictions rather than simply discarded. Across all benchmarks, Perseus substantially improved the PPV and LC-PPV, often approaching 0.99, while substantially reducing the FAR. While this reduces species-level sensitivity in some cases, it reflects the biological reality of many environmental microbiomes, where extensive taxonomic novelty and incomplete reference databases limit the reliability of fine-rank assignments. In such contexts, confident assignments at higher taxonomic levels are often more informative than spurious species-level predictions.

Performance improvements were most pronounced for long reads and assembled contigs, where spatial context allows more reliable discrimination between consistent lineage signal and localized conserved matches. Importantly, Perseus introduces only modest computational overhead and operates directly on standard Kraken2 outputs without requiring modification of reference databases or classifier internals. This design allows Perseus to be easily integrated into existing Kraken2-based workflows for large-scale metagenomic studies. More broadly, this work suggests that frameworks for taxonomic classification of long reads or assembled contigs benefit from separating evidence accumulation from confidence estimation. Existing k-mer classifiers are highly effective at aggregating sequence evidence, but they do not explicitly evaluate whether that evidence forms a spatially and hierarchically consistent pattern along the sequence. By reframing taxonomic classification as a lineage-aware confidence estimation problem, Perseus identifies the deepest taxonomic rank supported by coherent evidence rather than forcing a single-rank prediction.

Beyond our current implementation, several research opportunities remain. Perseus does not attempt to reclassify sequences beyond the lineage proposed by Kraken2 and therefore cannot assign precise labels to taxa that fall outside the inferred lineage. Our experiments show that this is predominantly due to gaps in the reference databases rather than limitations of the Perseus model, although future work could extend this framework to estimate the degree of taxonomic novelty relative to the reference database. This may help guide downstream discovery workflows such as targeted assembly of previously uncharacterized lineages. In addition, extending lineage-aware confidence estimation to operate on outputs from additional classifiers, such as Centrifuger or Metabuli, represents a promising direction for classifier-agnostic taxonomic refinement. Finally, our evaluation has been focused on bacterial genomes. Extending Perseus to archaeal, viral, and eukaryotic sequences represents an important step for future work.

In summary, Perseus provides a scalable and principled approach for reducing over-specific taxonomic assignments in long-read metagenomics. By modeling spatial and lineage-consistent patterns of k-mer evidence, Perseus produces classifications that more accurately reflect the strength and limits of the underlying sequence evidence, particularly in complex and under-characterized microbial communities. This demonstrates how separating evidence accumulation from confidence estimation improves robustness in long-read metagenomic classification.

## Code availability

The code for Perseus is available open-source on Github: https://github.com/matnguyen/perseus. Perseus can be installed through Conda, pip, or Docker. Scripts for reproducibility are available on Github: https://github.com/matnguyen/perseus-scripts.

## Acknowledgments

We would like to thank our collaborators in the BioDIGS consortium. We would also like to thank Ben Langmead, Steven Salzberg, Sina Majidian, Omar Ahmed, Mahler Revsine, Vikram Shivakumar, Alexander Sweeten, and Alex Ostrovsky for helpful discussions.

## Supplementary Material

**Figure S1:**
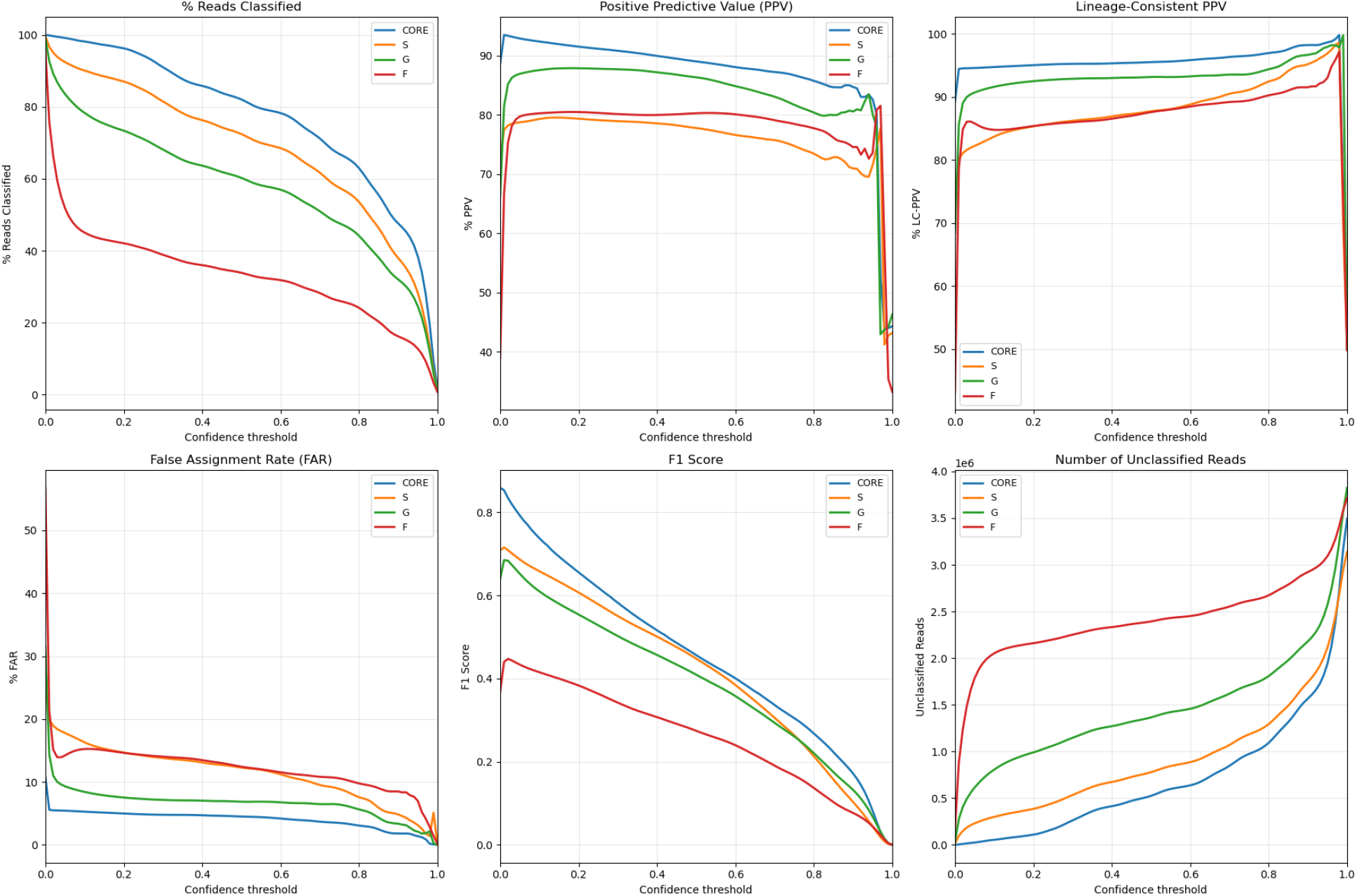
Alignment performance and query coverage as a function of rank back-off from Kraken to Perseus on the German forest microbiome. Bars show the proportion of reads that successfully aligned to the reference (alignment rate on the left y-axis), while the overlaid line shows the median query coverage among aligned reads (right y-axis). Categories on the x-axis denote the original Kraken assignment at species level and the corresponding Perseus back-off rank (species through superkingdom). As classifications are progressively backed off to higher taxonomic ranks, both alignment rate and median query coverage decrease.

**Figure S2:**
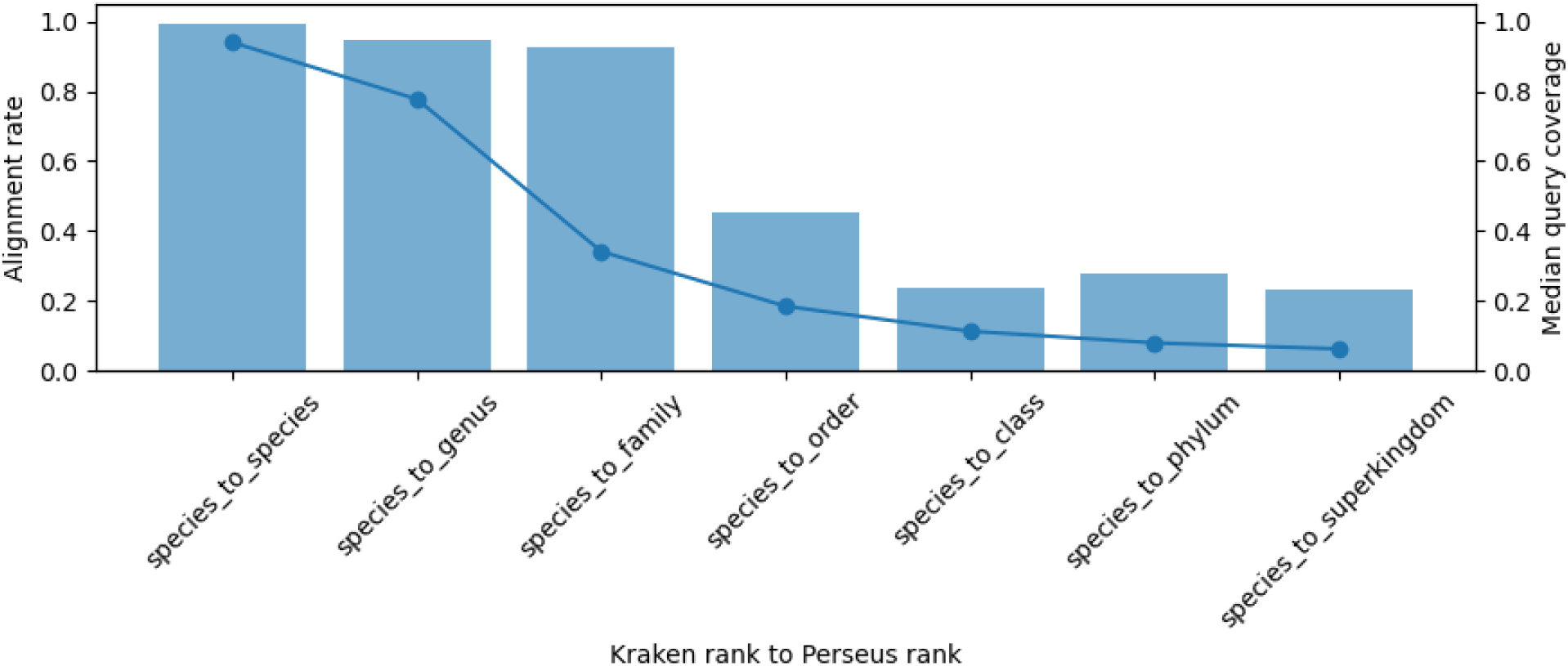
Performance of Kraken2 across confidence thresholds in the inclusion/exclusion evaluation datasets. Metrics shown include (top row) percent reads classified, positive predictive value (PPV), and lineage-consistent PPV; (bottom row) false assignment rate (FAR), F1 score, and number of unclassified reads. CORE denotes the inclusion dataset; S, G, and F denote species-, genus-, and family-exclusion datasets. As confidence increases, call rate decreases while precision improves, illustrating the sensitivity–precision tradeoff under increasing taxonomic novelty.

